# Trans-generational maintenance of mitochondrial DNA integrity in oocytes during early folliculogenesis

**DOI:** 10.1101/2025.01.02.631059

**Authors:** Qin Xie, Haibo Wu, Jiaxin Qiu, Junbo Liu, Qifeng Lyu, Hui Long, Wenzhi Li, Shuo Zhang, Xueyi Jiang, Yuxiao Zhou, Yining Gao, Aaron J. W. Hsueh, Yanping Kuang, Lun Suo

## Abstract

Mutations in mitochondrial DNA (mtDNA) can lead to mitochondrial and cellular dysfunction. However, recent studies suggest that there is purifying selection against mutant mtDNAs during transgenerational transmission. We investigated the mtDNA dynamics during ovarian follicle development. Using base-editing, we generated mice harboring a 3177 G>A mutation corresponding to the human Leber hereditary optic neuropathy (LHON)-related mtDNA mutation and confirmed the transgenerational reduction of mutant mtDNA. Using a mouse follicle culture system in which pathogenic mtDNA mutations were introduced *in vitro* followed by mtDNA sequencing and digital PCR, we found this germline heteroplasmy shift during early folliculogenesis was induced by the decrease of mutant mtDNA together with compensatory replication of wild-type mtDNA. In contrast, synonymous mtDNA mutations did not impact the mtDNA dynamics. These findings show that mice can eliminate certain pathogenic mtDNA mutations in the germline during early folliculogenesis, maintaining mitochondrial integrity across generations.

## Introduction

The mitochondrial genome (mtDNA) is a circular, double-stranded DNA located within the mitochondrial matrix and encodes 37 genes critical for oxidative phosphorylation^1^. There are usually thousands of copies of mtDNA in each somatic cell but more than 100,000 copies in a mature oocyte. Due to the strong exposure to oxygen free radicals, inadequate proofreading and repair mechanisms, and strict maternal inheritance pattern^2^, the mtDNA, unlike the nuclear genome, is highly susceptible to the accumulation of pathogenic mutations^3,4^. The proportion of mutations (heteroplasmy) in crucial areas of mtDNA above a threshold will lead to serious diseases such as human Leber hereditary optic neuropathy (LHON), mitochondrial encephalomyopathy, lactic acidosis and stroke-like episodes (MELAS)^5,6^.

Recent studies suggest that mutant mtDNAs are decreased during transgenerational transmission. The elimination of pathogenic mutations across generations, also called germline purifying selection, was first observed in PolgA mutant mice^7^ and mice with a mutation in ND6^8^. This phenomenon has also been found in Drosophila *melanogaste*r^9–12^ and nematodes^13^. Indeed, mother-offspring pairs in humans also showed similar elimination of pathogenic mutations across generations based on large-scale observational data^14–17^.

Purifying selection is of paramount importance for the adaptation and propagation of populations while avoiding extinction, but the mechanisms are still poorly understood. To be transmitted to offspring, the mtDNA first segregates into primordial germ cells (PGC) and then replicates during prolonged folliculogenesis^18,19^. However, the difficulty of culturing germ cells *in vitro* and manipulating mtDNA mutations has limited progress in this area. Here, we employed a novel mtDNA editing tool^20^ to introduce a 3177 G>A mutation of mtDNA (m. 3177G>A) in mice corresponding to the human Leber hereditary optic neuropathy^21^. Utilizing this model and introducing mutations into follicles cultured *in vitro* by a recently developed culture system^22–27^, we investigate the dynamics of mtDNA purifying selection during folliculogenesis. In this work, we first provide convincing evidence that the pathogenic but not synonymous mtDNA mutations could be eliminated during folliculogenesis. Secondly, it has been observed that selection against mtDNA mutations occurs at the organelle level by two pathways, by which compensatory replication of wild-type mtDNA copies and elimination of mutant mtDNA copies work together within one oocyte. Moreover, the experimental platform developed by this work could also serve as a direct tool to study the molecular mechanism of purifying selection in follicles.

## Materials and Method

### Mice

Mice were purchased from Beijing Vital River Laboratory Animal Technology Co., Ltd. and kept at 23°C with a 12h:12h light-dark cycle. The Institutional Animal Care or Research Ethics Committee at Shanghai Jiao Tong University School of Medicine approved all experimental procedures.

### Plasmids generation

The TALE-based DdCBEs vectors are composed of mitochondrial localization sequence (MTS), N terminal, C terminal, one of four split DddA halves, and UGI-coding sequences20. The Talen RVDs were assembled by the Golden-Talen system. Table S1 lists all binding array of each DdCBE element.

### In vitro Transcription

TALE-based DdCBEs plasmids with T7 promoters were linearized with NotI (NEB) endonuclease and purified using 1.2% electrophoresis gel. Based on the manufacturer’s manual, the purified product was employed as the template for in vitro transcription (IVT) using the mMESSAGE T7 ULTRA kit from Life Technologies. To prepare the TALENs mRNA for injection into germ cells, both halves were purified using a MEGAclear kit from Life Technologies.

### Construction of mtDNA mutant animals

To construct animal model with the m.3177 G>A mutation, six-week-old female mice (C57/B6) were super-ovulated for the collection of zygotes using intraperitoneal injections of 10 IU of PMSG, followed by 10 IU of hCG (Ningbo Hormone Products Co., China) 48 hours later. Then, male and female mice were allowed to mate. Twenty hours after hCG treatment, fertilized eggs with two pronuclei were removed from oviducts and placed in M2 medium. Cumulus mass was removed in 70μg/mL hyaluronidase for microinjection. After the pronuclear embryos had recovered for an hour, forward and reverse TALENs-based DdCBE-3177 were mixed and injected into the cytoplasm of fertilized eggs, followed by culturing in the KSOM media at 37°C under 5% CO2 in air under the mineral oil. For generating the mice model, two-cell stage embryos were transferred in to the fallopian tube of pseudo-pregnant mice.

### Follicle culture in vitro

A detailed method has been described previously22-26. Briefly, 12-day-old B6D2F1 females were used to collect secondary follicles with a diameter between 105 and 125μm. Isolated follicles with intact shape and high density of granulosa cells were chosen for further microinjection and in vitro culture. Follicles were microinjected with a mixture of the left and right halves of DdCBE mRNA after recovering for an hour.

Following microinjection, a collagenase treatment lasting 25 minutes was carried out to dissolve the follicular wall. Follicles were grown in α-MEM with 5% fetal bovine serum, 1% GlutaMax (Gibco), 2% PVP, 0.1% Penicillin-Streptomycin (Gibco), 150M 2-O-alpha-d-glucopyranosyl-l-ascorbic acid (AA2G, Tokyo Chemical Industry), and 100mU/mL follicle-stimulating hormone (FSH, Gonal-F). The incubator was set at 37°C, 100% humidity, and 5% CO2 in the air, and half of the media was changed every other day.

### DNA extraction

The extraction of genomic DNA from mouse tissues was performed using the DNeasy Blood & Tissue kit (Qiagen). To extract the DNA from single oocyte, follicles were subjected to digestion using a combination of 0.5% trypsin (Gibco), 0.1% collagenase Type I (Worthington), 0.1% DNase I (Sigma), and 1mM EDTA (Solarbio) for 10 minutes. Subsequently, oocytes were stripped of granulosa cells before three rinsings in a solution containing 0.4% BSA. Subsequently, they were immersed in 5μl of QuickExtract™ DNA Extraction Solution for digestion.

### Digital PCR

The primers and probes were purchased from IDT, as shown in Table S2. A 5μl sample containing a single oocyte was added with 10μl of DNase-free ddH2O. Subsequently, a 2μl portion of this mixture was utilized as a template for PCR amplification. The reaction droplets were generated using the droplet generator of the QX200TM droplet digital PCR Instrument from a PCR mix of 20μl.

Subsequently, the droplets were amplified on the Bio-Rad T100 PCR instrument using the following cycling conditions: an initial denaturation step at 95°C for 10 minutes, followed by 40 cycles of denaturation at 94°C for 30 seconds, annealing at 60°C for 1 minute, and extension at 72°C for 1 minute. The final extension step was performed at 98°C for 10 minutes. Following PCR amplification, the fluorescence signal emitted by FAM (representing wild-type copies) and HEX (representing mutant copies) within each droplet was detected using the QX200 Droplet Reader and subsequently analyzed using the QuantaSoft software.

### Target sequencing

A quantity of 100ng of genomic DNA obtained from tissues or lysates of single cells was utilized for the initial PCR amplification. Phanta Flash DNA Polymerase (Vazyme), together with primers that incorporated barcodes and Illumina adapters (Table S2) were used to amplify the desired target sequences. A volume of 1 microliter of the product was utilized for the second-round PCR employing index primers from Vazyme. Following the completion of the second-round PCR, the samples containing distinct barcodes and indexes were combined and subjected to purification through gel extraction utilizing the QIAquick Gel Extraction Kit manufactured by Qiagen. Subsequently, the quantification of the purified samples was performed using the Qubit ssDNA HS Assay Kit provided by Thermo Fisher Scientific.

The Illumina NovaSeq 6000 platform was utilized for deep sequencing. The sequencing data underwent quality control monitoring using fastp (v0.23.2). The sequencing reads were demultiplexed using fastq-multx (v1.4.1) software with the barcoded PCR primers. Subsequently, the mutation rates of the on-target sites were determined by analyzing the output file obtained from batch analysis using CRISPResso2 (v2.0.32). Statistical analysis was performed using custom scripts developed in R (v4.2.1)

### Whole mtDNA sequencing

Whole mitochondrial DNA (mtDNA) was amplified using long-range PCR, which involved amplifying two overlapping 8kb fragments. The primer sequences used are listed in Table S2. The PCR products underwent purification using the QIAquick Gel Extraction Kit (Qiagen) and were subsequently utilized as input for library construction using the TruePrepTM DNA Library Prep Kit V2 for Illumina (Vazyme). Prior to conducting deep sequencing, the libraries underwent purification using DNA clean beads and were subsequently quantified using the Qubit ssDNA HS Assay Kit from Thermo Fisher Scientific.

To analyze next generation sequencing data from whole mitochondrial genome sequencing, the initial step involved aligning the qualified reads to the mitochondrial reference genome (mm10) using BWA (v0.7.12) with the mem – M. Subsequently, BAM files were generated using SAMtools (v.1.9). Positions with conversion rates ≥ 1% were identified among all cytosines and guanines in the mitochondrial genome using the REDItoolDenovo.py script from REDItools (v.1.2.1).

### Transmission electron microscope observation

The specimen underwent a series of preparation steps for transmission electron microscope observation. Initially, it was immersed in a solution consisting of 2.5% glutaraldehyde and 2% paraform for 2 hours at 4℃. Following this, the specimen underwent two washes in phosphate buffer and was subsequently fixed using 1% osmium tetroxide.

Dehydration was carried out through sequential immersions in increasing concentrations of ethanol (30%, 50%, 70%, 80%, 95%, and 100%) for approximately 10 minutes each. Further processing involved two changes of 100% propylene oxide (P.O.) for 15 minutes per change. Subsequently, it was placed in a mixture of P.O. and the embedding resin (epon 812) in varying ratios for the required duration, followed by immersion in pure embedding resin (epon 812) for 6 hours at 37℃.

The specimen was then encapsulated in embedding medium and subjected to heating at 60℃ for 48 hours. Subsequent steps involved sectioning the samples to 70-90nm using a diamond knife (Leica EM UC7), followed by staining with uranyl acetate and lead citrate before observation under a transmission electron microscope (HITACHI H-7650 or FEI Talos 120).

### Quantification and statistical analysis

For method of mapping the density curve, we first generated the density curve of the observed heteroplasmy shift. Then we generated 1000 times the sample size random value based on the Kimura distribution and drew the density curve. For data analysis, the normality of the data was evaluated through the Shapiro-Wilk test. If the data satisfied the assumptions of normality, we employed an unpaired Student’s t-test to examine the differences between groups. If the data failed to satisfy the assumptions of normality or equal variances, we utilized the Mann-Whitney U test. Statistical analysis was performed by GraphPad Prism 9 and P<0.05 will be considered as significant. The statistical details could be found in each figure legend.

## Results

### The pathogenic m.3177G>A mutation was eliminated in mice during germline transmission

To explore the germline transmission of mutant mtDNA, we first constructed a mouse model with a m.3177 G>A mutation (Figure 1A and 1B) using a recently developed mtDNA editing tool DdCBE^20^, and the detailed information of the DdCBE is shown in Table S1. *In vitro* transcribed 3177-DdCBE mRNA was injected into zygotes to generate the F0 mice. These F0 females were then backcrossed with wild-type (WT) males. After screening F1 mice, one female carrying a high ratio of m.3177 G>A mutation without off-target mutations in mtDNA (Figure 1A and 1C) was selected as the founder of the m.3177 G>A mouse model (Figure 1A). Sequencing of mtDNA in 17 different tissues indicated stable heteroplasmy and the average coefficient of variation was 0.077% (Figure 1D). We then used tail sequencing data to monitor heteroplasmy during subsequent generations. To observe the purifying selection of mtDNA mutation, we bred a total of 44 mother and their 793 pups from F1 to F5 generations. As expected, the pups’ heteroplasmy exhibited a wide variation likely due to the bottleneck effect^18,28^. Further analysis of heteroplasmy in mother-pup pairs revealed a strong negative selection of m.3177 G>A mutation, as evidenced by a slope of 0.69 on the mother-offspring fitness curve (*P* < 0.0001) (Figure 2A). To analyze the relative ratio of the heteroplasmy to the mother, we transformed the heteroplasmy by ln(h(1-h0)/h0(1-h)). The variable “h” is the pup’s heteroplasmy, and “h0” represents the mother’s heteroplasmy (Figure 2B). Both analyses showed that the average heteroplasmy in pups was consistently lower than the mother’s heteroplasmy, suggesting purifying selection of pathogenic mutation during trans-generation transmission. We also took advantage of another mouse model with a synonymous mutation due to the off-targeting of DdCBE. By contrast, mice carrying a synonymous mutation m.6134 C>T showed an unbiased transmission (Figure 2C-2D), indicating no purifying selection.

**Figure 1.**
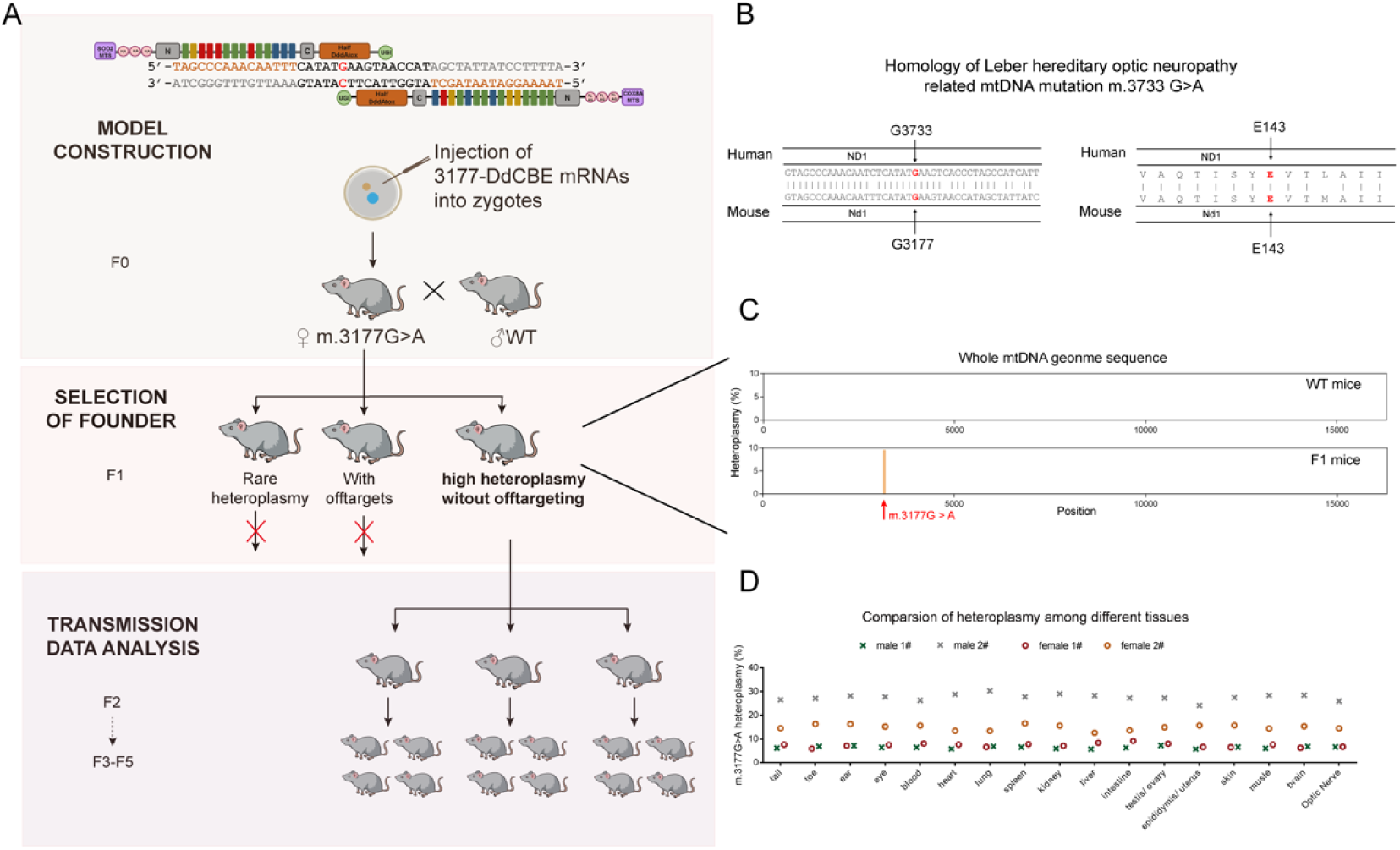
Generation of m.3177 G>A mutant mice. (A) The experimental workflow for generating the m.3177 G>A mice by injecting base editing tool-DdCBE-3177 mRNAs-into zygotes and selection of high heteroplasmy lines without off-targeting. (B) Homology of the human Leber hereditary optic neuropathy mtDNA mutation m.3733 G>A and corresponding mouse mutant selected. (C) Lack of off-targeting in the mtDNA genome in the F1 m.3177 G>A mice. A wildtype (WT) mouse was used as the control. (D) The heteroplasmy among different tissues in m.3177 G>A mice.

**Figure 2.**
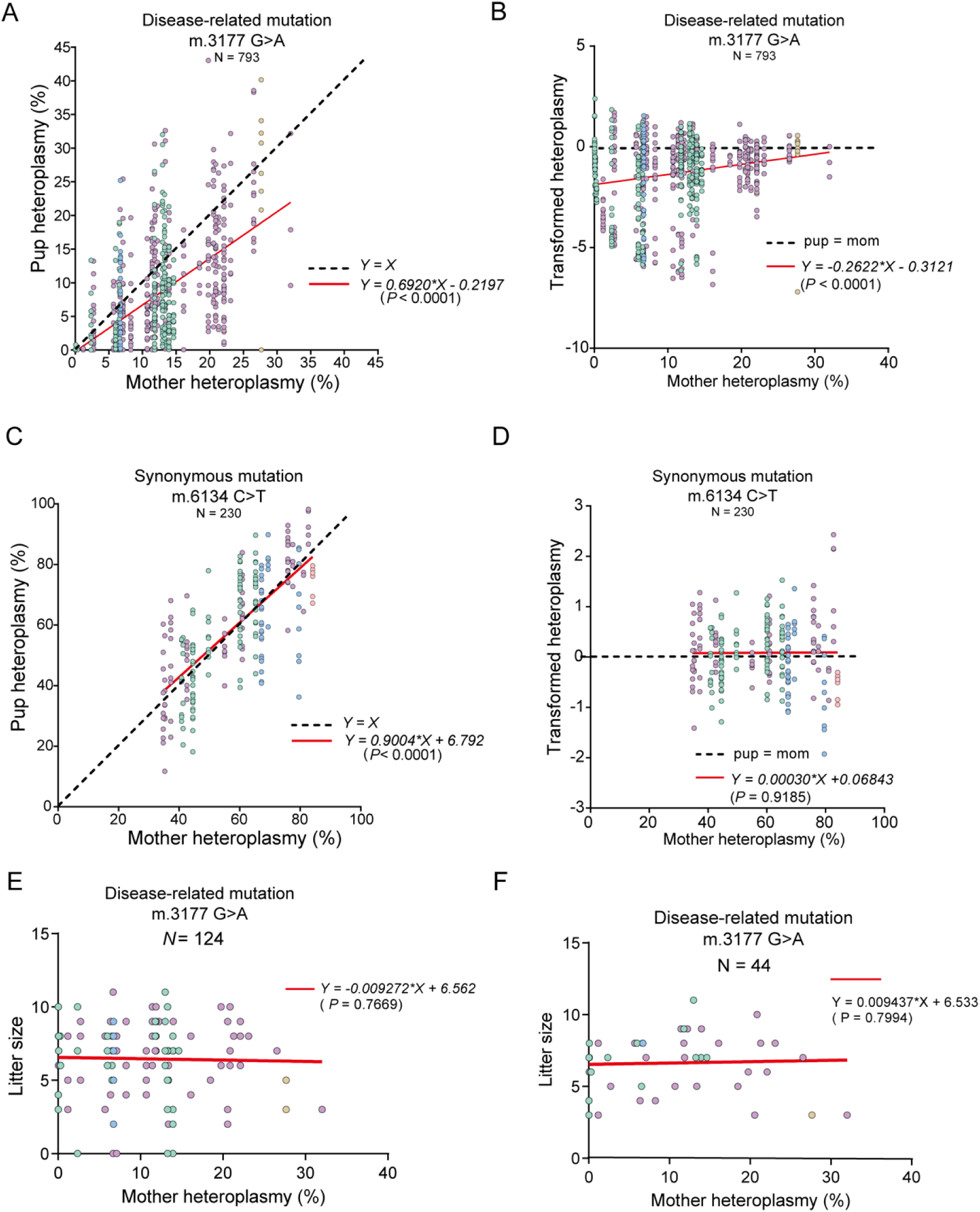
Germline transmission of mtDNA m.3177 G>A mutation in mice. (A)Transmission of heteroplasmy from 44 mothers to 793 pups in disease-related m.3177 G>A mutant mice. (B)The transformed heteroplasmy shift of disease-related m.3177 G>A mutant mice shown in Figure 2A. (C)Transmission of heteroplasmy from 20 mothers to 230 pups of a synonymous mutation in m.6134 C>T mice. (D)The transformed heteroplasmy shift of synonymous mutation with m.6134 C>T mice shown in Figure 2C. (E)The litter sizes of 124 deliveries in m.3177G>A mice. (F)The litter size at 44 first deliveries of m.3177G>A mice. The transformed heteroplasmy shift was calculated as ln(*h*(1-*h_0_*)/*h_0_*(1-*h*)). The variable “*h”* is the pup’s heteroplasmy, and “*h_0_*” represents the mother’s heteroplasmy. The red line represents the fitted curve while the dashed line indicates unbiased transmission. For panels A-F, orange, blue, green, purple, and yellow dots refer to F1, F2, F3, F4, and F5 generations, respectively.

To test if this heteroplasmy decline was derived from embryonic lethality with high heteroplasmy, we analyzed the relationship between litter size and mother’s heteroplasmy ratios in m.3177 G>A mutant mice and found no correlation (Figure 2E, *P* = 0.7669). To avoid the confounder of maternal age on litter size, subgroup analysis focusing on the first litter of each mother was performed and similar results were obtained (Figure 2F, *P* > 0.05).

### Elimination of the mtDNA mutation with m.3177G>A occurs in early folliculogenesis

The decreased mutation load observed in offspring without affecting litter sizes implied that selection pressure could act before ovulation, consistent with previous studies^8,29,30^. Before maturation for fertilization, the oocytes in primordial follicles pass through primary, early secondary, late secondary, and antral stages (Figure 3A). A previous study reported no selection of m.3875delC in primordial germ cells and P3.5 oocytes (follicle starting to assemble), indicating selection occurred after follicle formation^29^. In contrast, another study found that the m.5024 C>T mutation in a mtDNA tRNA gene decreased in primary and secondary follicles but increased in antral follicles^31^. To determine the stage of purifying selection during folliculogenesis, we isolated different stages of follicles from the ovaries of four mice with the m.3177 G>A mutation followed by sequencing mtDNAs of the oocytes. To combine the data of mice with various levels of heteroplasmy, we normalized absolute heteroplasmy into heteroplasmy shift in pups as compared to the respective mother. In primary follicles, the heteroplasmy exhibited a slight decrease from 1 to 0.93 (Figure 3B-3C). With further development, the heteroplasmy decreased substantially to 0.79 in the early secondary follicle stage and further to 0.74 in late secondary follicles, a level comparable to those found in offspring (0.70) (Figure 3B and 3D-3F). Hence, a clear decrease in heteroplasmy in pups was mainly occurred during early folliculogenesis.

**Figure 3.**
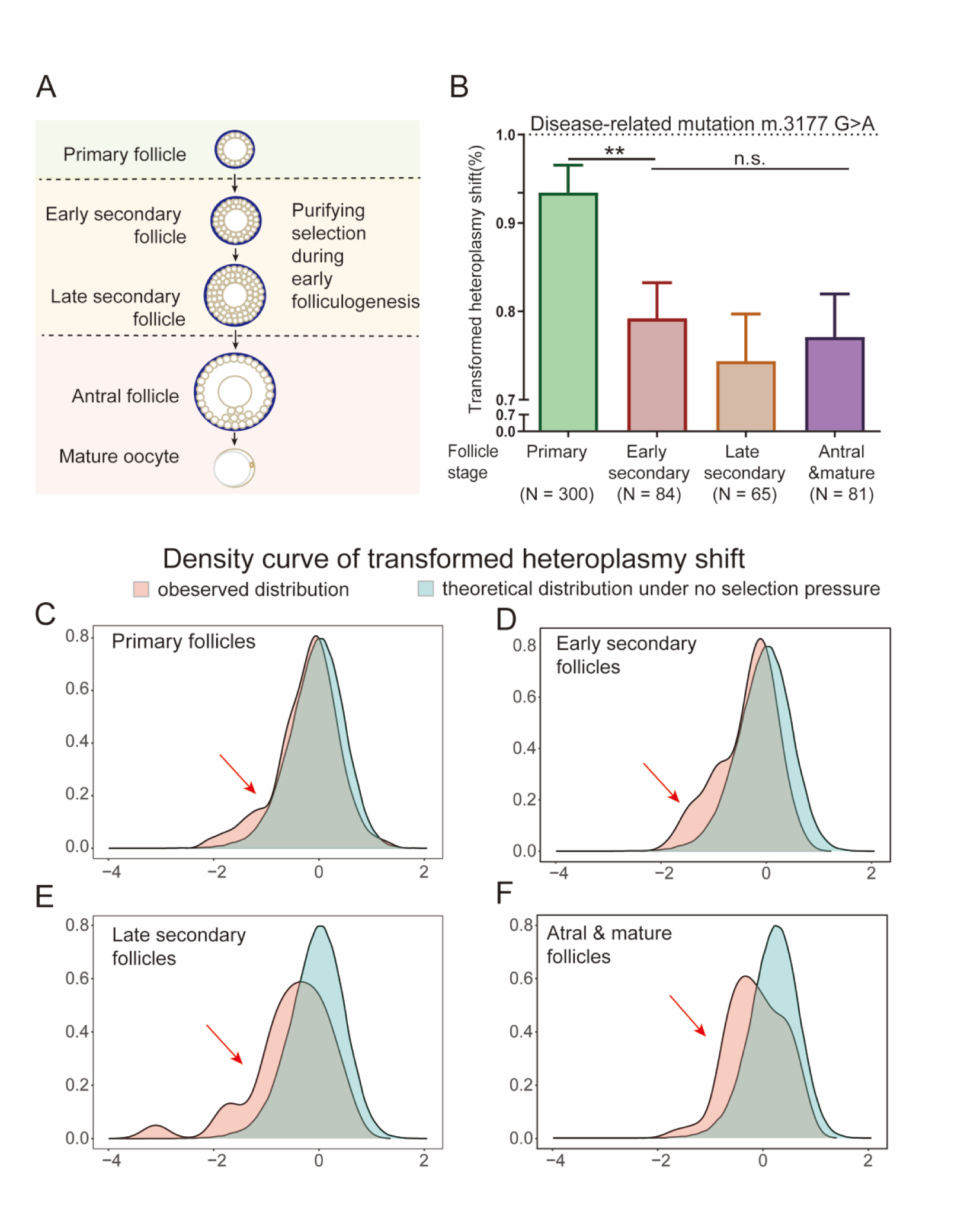
mtDNA dynamics in oocytes carrying the m. 3177 G>A mutation. (A)The diagram of follicle development. (B)The transformed heteroplasmy shift of mtDNA in oocytes at different follicle stages in m.3177 G>A mice showing a drop in heteroplasmy during the transition from primary to secondary follicle development. The heteroplasmy shift was calculated as *h*(1-*h_0_*)/*h_0_*(1-*h*). *h* is the oocyte heteroplasmy, and *h_0_* is the mother’s heteroplasmy. Significance was calculated with *Mann-Whitney U* test (** *P* < 0.01). (C-F) Density curve of transformed heteroplasmy shift in primary follicles (C), early secondary follicles (D), late secondary follicles (E), antral and mature follicles (F).

### Follicular elimination of the mtDNA mutations in a functional dependent manner

To further investigate the dynamics of mitochondrial mutations during folliculogenesis, we isolated early secondary follicles from WT mice based on a recently reported culture system^22–27^. The follicles cultured *in vitro* exhibited similar morphology to those grown *in vivo*, and 50% of them could reach the blastocyst stage^22,23^, indicating the developmental potential of *in vitro* cultured follicles (Figure 4).

**Figure 4.**
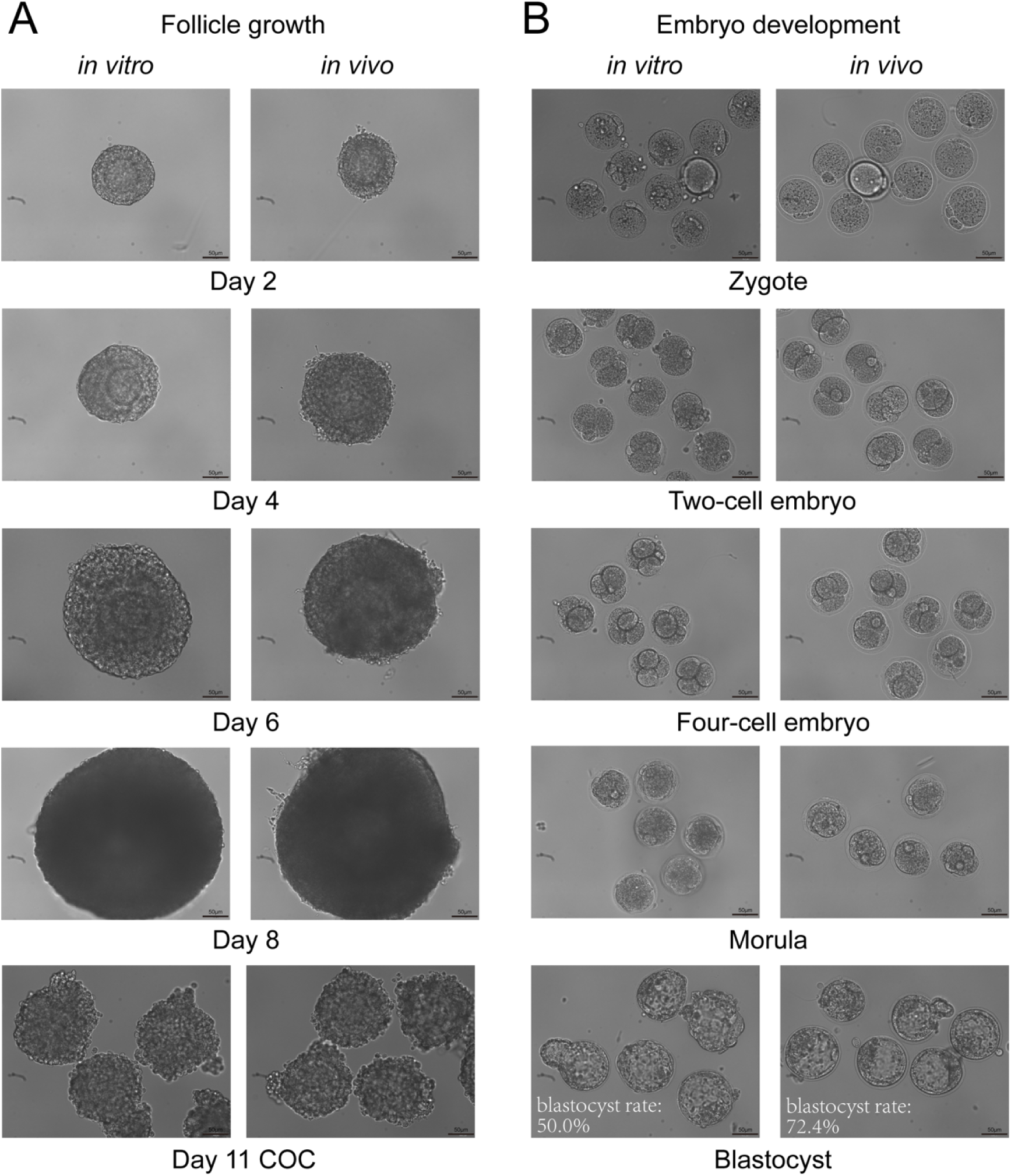
The *in vitro* development of secondary follicles. (A) Comparison of the morphology of secondary follicles derived from *in vivo* and *in vitro* models. (B) Comparison of the morphology of embryos derived from secondary follicles derived *in vivo* and *in vitro*.

We injected the DdCBE-3177 mRNA into oocytes of early secondary follicles on the day of isolation (Day 0) and sampled them continuously for seven days (Figure 5A). One day after microinjection, the DdCBE-3177 introduced about 30% mutations. This was followed by a sharp drop by day 4 of culture observed in three independent experiments (Figure 5B). We also generated another disease-related mutation m.12918 G>A, corresponding to the human m.13513 G>A mutation causing Leigh Syndrome and MELAS^32,33^. A similar decline of heteroplasmy was observed in culture (Figure 5C).

**Figure 5.**
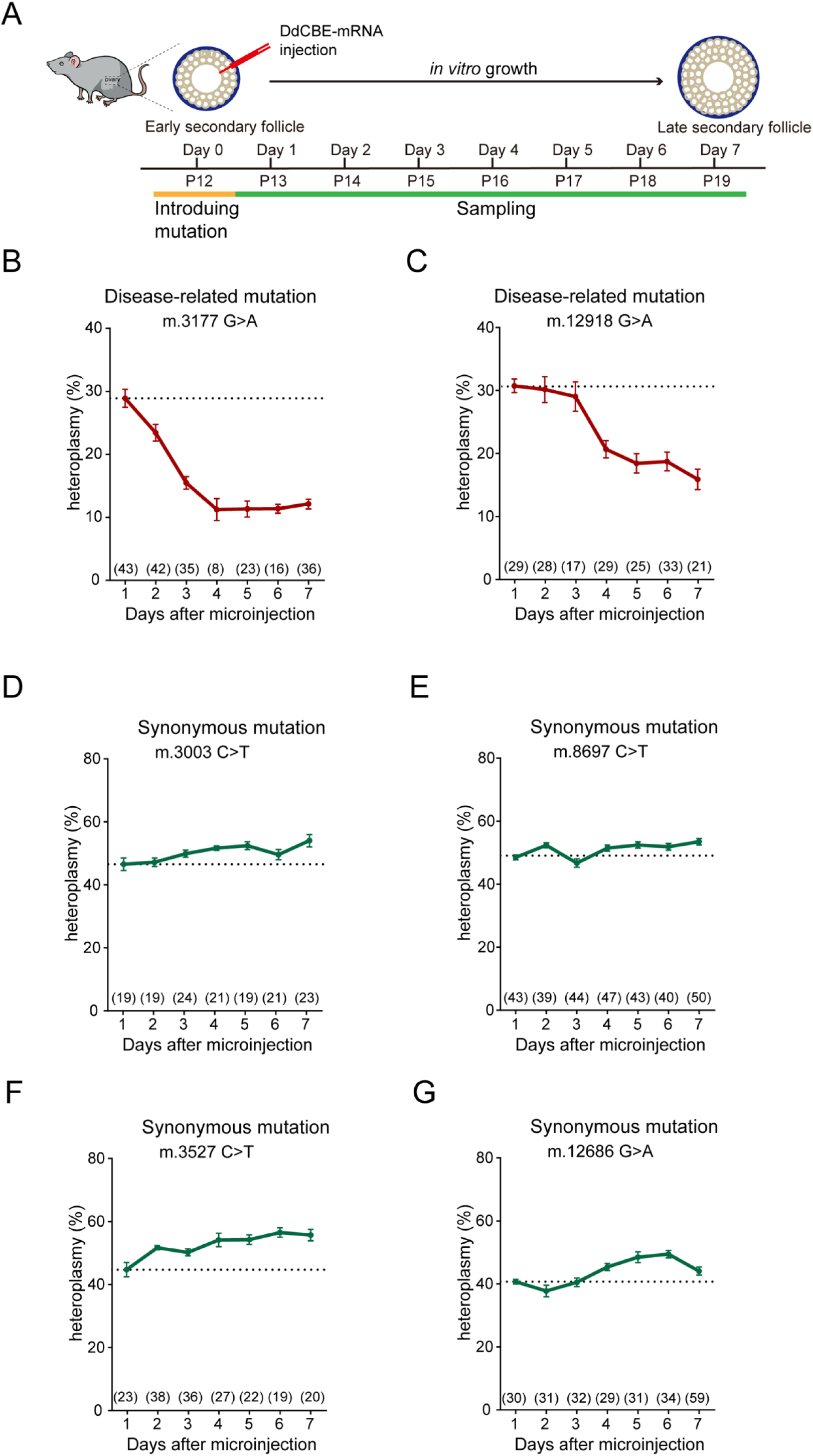
Dynamic of mtDNA disease-related and synonymous mutation rates during early folliculogenesis in *vitro*. (A) *In vitro* culture and microinjection of DdCBE-mRNAs for base-editing and sampling at different times during secondary follicle development. P12 to 19 denote corresponding to postnatal days *in vivo*. (B-C) Dynamics of disease-related mtDNA heteroplasmy for m.3177 G>A (B) and m.12918 G>A (C) mutations during secondary follicle development *in vitro*. (D-G) Dynamics of synonymous mtDNA heteroplasmy for m.3003 C>T (D), m.8697 C>T(E), m.3527 C>T (F), and m.12686 G>A (G) during secondary follicle development *in vitro*. The error bars represent the standard error of the mean (SEM). Data was derived from 3 independent replicate experiments. The number in parentheses indicates the number of oocytes sequenced.

To discern if the purifying selection was directed against base substitutions or dysfunctional mitochondria associated with pathogenic mutations, we introduced four synonymous mutations (m.3003 C>T, m.8697 C>T, m.3527 C>T, and m.12686 G>A) that altered mtDNA bases while preserving encoded proteins. Unlike the disease-related mutations, we observed no purifying selection of these synonymous mutations during folliculogenesis (Figure 5D-5G). Thus, selection pressure might arise from functional impairment of mitochondria caused by pathogenic mutations.

### Heteroplasmy decrease of the m.3177G>A mutation in oocytes was induced by eliminating of mutant copies together with compensatory replication of wildtype variants

It is unclear whether decreases of heteroplasmy are the result of the elimination of mutant copies, the compensatory upregulation of WT copies, or both^4,34,35^. To investigate the molecular dynamics during selection, we employed digital PCR on oocytes in cultured follicles to quantitate copy numbers for both WT and mutant DNAs in individual oocytes. We first examined the dynamics of mtDNA copy numbers of unedited oocytes over seven days of culture and found a rapid increase in mtDNA copy numbers to approximately 100,000 copies within the first three days and remained stable later (Figure 6A). After introducing the pathogenic m.3177 G>A mutation into oocytes of secondary follicles, we found a decline in mutant copy numbers, accompanied by an increase in WT copy numbers during the seven days of culture (Figure 6B). To determine if the decrease of mutant copies is due to mtDNA degradation or mitophagy, we performed transmission electron microscopy (TEM) of oocytes. Compared with the mitochondria with normal morphology in unedited oocytes, the mitochondria in edited oocytes were swollen and located in the lysosome (Figure 6E-6G), suggesting the activated mitophagy. Thus, the observed decrease of pathogenic mutations was due to both elimination of mutant copies and compensatory upregulation of WT copies.

**Figure 6.**
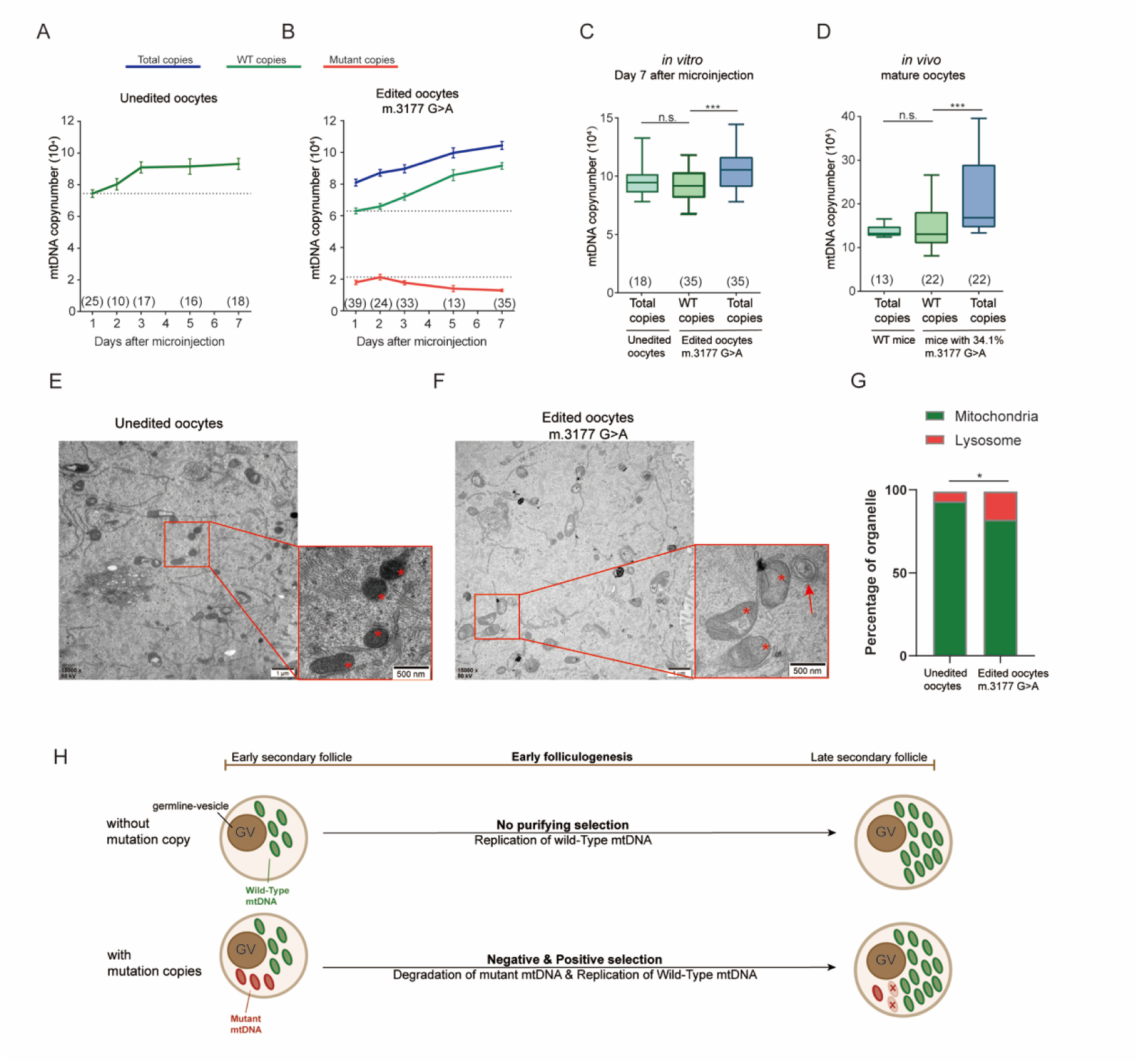
Dynamic of mtDNA copy numbers changes during early folliculogenesis. (A) The dynamic of mtDNA copy numbers derived from digital DNA PCR in unedited oocytes during secondary follicle development *in vitro*. The green line represents wild-type (total) copies. (B) The dynamic of mtDNA copy numbers in m.3177 G>A edited oocytes during *in vitro culture*. The blue line represents the total copies. The green line represents wild-type copies. The red line represents mutant copies. The error bars represent the standard error of the mean (SEM). Data was derived from 3 independent replicate experiments. (C) Comparison of mtDNA copy numbers in edited oocytes and unedited oocytes on Day 7 after microinjection. Significance was based on two-tailed *Student’s t-test* (*** p<0.001). The data are presented by box plots (maximum-median-minimum). (D) Comparison of mtDNA copy numbers in mature oocytes from m.3177G>A and wildtype (WT) mice. Significance was calculated based on the *Mann-Whitney U* test (*** p<0.001). The data are presented by box plots (maximum-median-minimum). (E) Transmission electron microscope images of mitochondria in unedited oocytes. The red asterisk points indicate the mitochondria. (F) Transmission electron microscope images of mitochondria in m.3177 G>A edited oocytes. The red asterisk points indicate the swollen mitochondria and the red arrow indicate the lysosome. (G) Comparison of the ratio of mitochondria and lysosome of unedited (8 independent slides) and edited (13 independent slides) oocytes from transmission electron microscope images. Significance was based on Chi-squared *test* (*p<0.05). (H) Diagram showing dynamics of mtDNA copy number changes during early folliculogenesis. During early folliculogenesis, unedited follicles showed replication of wildtype (WT) copies while edited follicles showed simultaneously negative selection by degradation of mutant copies and compensatory replication of WT copies.

Further analysis of digital PCR data from both *in vitro* and *in vivo* models revealed that mutant and WT oocytes gained comparable copies of WT mtDNA during folliculogenesis. On Day 7 of culture, the WT copy numbers in edited oocytes had risen to a comparable level to those in unedited oocytes, with the total copy numbers in edited oocytes exceeding those in unedited oocytes (Figure 6C). We also performed digital PCR for mature oocytes obtained from mature follicles of mutant and WT mice pretreated with gonadotropins. Similarly, the WT copy numbers of *in vivo* matured oocytes from m.3177 G>A mice were also comparable to those from WT mice (Figure 6D), although the *in vivo* oocytes from mutant mice exhibited a wide range of heteroplasmy and copy number of mtDNA. Thus, during early folliculogenesis, oocytes in unedited follicles showed replication of WT copies while those in edited follicles showed both degrading mutant copies and replicating WT copies to reach comparable copy numbers (Figure 6H).

## Discussion

Mitochondria are the energy factory of cells, possessing their own genome-mtDNA^1^. Pathogenic mutations on the mtDNA can lead to severe diseases, affecting multiple tissues, including the heart, brain, and muscles. Therefore, one of the key issues to maintaining human health is how to prevent the spread and accumulation of pathogenic mutations through generations^5,6^.

The Muller’s ratchet, first proposed by American geneticist Hermann Joseph Muller, indicates that in asexual reproduction without chromosome recombination (such as strict maternal inheritance of mtDNA^2^), mutations would accumulate continuously in offspring, gradually increasing until extinction of the population^3,4^. However, researchers have found in various species including humans^14–17^, mice^7,8^, Drosophila *melanogaste*r^9–12^, and nematodes^13^, that the pathogenic mutations are eliminated during trans-generational transmission, a process termed purifying selection. However, little is known about the mechanism currently.

In this study, by analyzing the heteroplasmy of oocytes at different follicle stages in the mouse ovary and observing the mtDNA dynamics in secondary follicles cultured *in vitro*, we provide evidence that purifying selection acts during folliculogenesis. From PGC to live pups, the germ cell goes through a long process of follicular development, oocyte maturation, fertilization, pre- and post-implantation embryo development. It holds important significance that purifying selection acts during folliculogenesis rather than other stages. Firstly, mtDNA in oocytes, unlike somatic cells, is amplified massively during folliculogenesis, and purifying selection can compensate for the lack of the repair mechanism and avoid mutation accumulation. Secondly, it avoids the dysfunction of mitochondria in important events such as oocyte maturation, ensuring optimal embryo development. Thirdly, purifying selection acting before ovulation safeguards offspring population size and reduces the risk of adverse events like stillbirths.

However, it is important to emphasize that different views exist in this field. In Mitalipov’s lab, Ma et al imply that purifying selection acts not in the maternal germline per se, but during post-implantation development ^36^. The reasons behind this difference in findings are worth further investigation. In our view, this may be due to the different models used between these two studies. The animals used in their study contained multiple mtDNA mutations which were caused by PolG deficiency rather than a single pathogenic mutation. As we know, there exists a well-known phenomenon of positive selection for some specific mutations, such as MELAS mutation m.3243A>G ^37,38^ or some other sites currently unknown. Therefore, a mixture of the multiple mtDNA mutations makes the purifying selection more complex, which might explain the difference between these two works.

Other than the timing for purifying selection, the level at which purifying selection occurs is also a critical problem in this field^35,39^. To decrease pathogenic mutations, purifying selection could act at four levels: genomic level (direct degradation or repair of mutation mtDNA), organelle level (mitophagy and/or preferential replication), cellular level (atresia of follicles or apoptosis of oocytes with high heteroplasmy), and organism level (lethality of pups with high heteroplasmy). First, the selection was unlikely to act at the genomic level, because of the unbiased transmission of synonymous mutations. Second, the increased ratio of swollen mitochondria in lysosomes found under TEM indicated selection at the mitochondrial level. Thirdly, purifying selection is unlikely at the cellular level because no follicle atresia has been found during early folliculogenesis until the early antral stage *in vivo*^40^ and the survival rate of edited oocytes is almost 100% during folliculogenesis. Lastly, the unaffected litter sizes of mice with the m.3177 G>A mutation excluded selection at the organism level. Thus, unlike the selection of nuclear genome mutations at the organism level, the selection of pathogenic mutations in the mitochondrial genome observed here likely occurred at the organelle level, due to mitochondrial demise.

Mitochondrial replacement therapy (MRT) is an important method for preventing the transmission of pathogenic mtDNA mutations to offspring, but it faces technical difficulties and ethical concerns regarding the ‘tri-parental’ construct^41^. Developing small-molecule drugs to eliminate pathogenic mutations before ovulation could offer novel reproductive options for women with mtDNA diseases, bypassing ethical concerns associated with MRT. Thus, exploring the underlying mechanism will pave the way to novel clinical therapies.

Our study pointed out two pathways of purifying selection: the selective elimination of mutant copies and the preferential replication of wild-type copies, ultimately obtaining an optimal number of wild-type copies. This replication bias made the wild-type mitochondria more likely to be inherited in the next generation. A study conducted in Drosophila *melanogaste*r also suggested the inhibition of replication of mutant mitochondria through inhibiting local translation^42^. In line with the direct decrease of mutant copies, previous studies indicated some mitophagy-related genes were involved in this process, including *Bcl2l13*, *ulk1*, *ulk2,* and *bnip3*^9,43^. Recently, the age effect has also been reported to affect the transmission of mtDNA ^44^, but the underlying mechanism is required to further study^35^.

Taken together, this study demonstrated the transgenerational elimination of pathogenic mutations during folliculogenesis through the degradation of mutant copies together with compensatory replication of WT copies, which advances our understanding of mitochondrial diseases and paves the way for novel therapeutic strategies.

## Acknowledgements

This work was supported by the Science and Technology Commission of Shanghai Municipality (21JC1403900 to L.S.), National Key R&D Program of China (2018YFC1003004 to L.S. and 2018YFC1003000 to Y.P.K.). The authors would like to express sincere gratitude to Jenny Yang from University of Cambridge for her generous help in revision of manuscript.

## Author contributions

L.S., Y.P.K., A.H., and Q.X. conceived the idea and designed the project. J.X.Q., Q.X., S.Z., X.Y.J., Y.X.Z. and Y.N.G. designed and generated the DdCBE plasmids. Q.X. performed animal and follicle experiments with the help of W.Z.L., Q.F.L. and H.L. Moreover, H.B.W., J.B.L. and Q.X. performed the microinjection. H.B.W. performed the Illumina sequence data analysis. Q.X. wrote the manuscript with the help of A.H. and L.S.

## Supplemental information

**Table S1.**
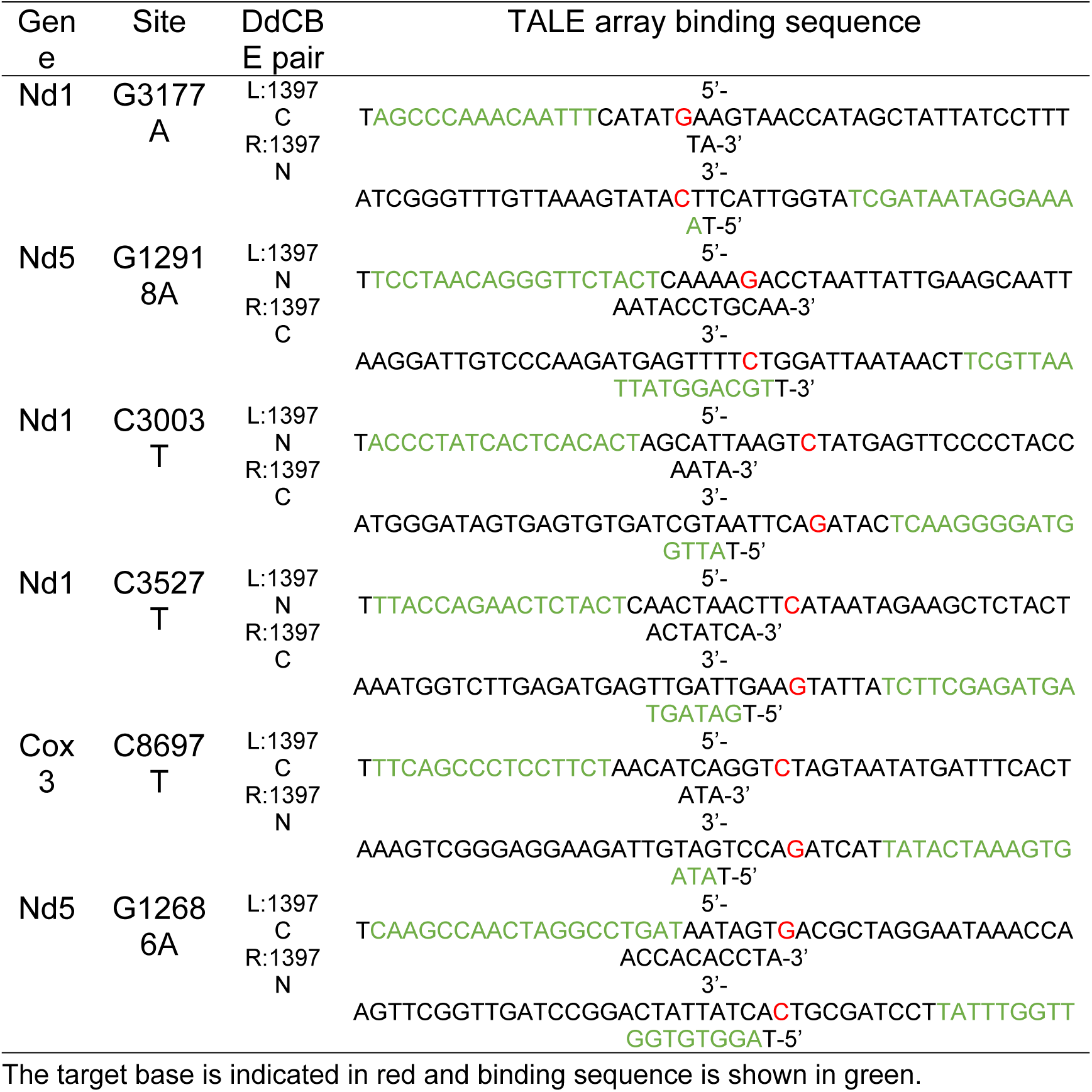
TALE array binding sequence.

**Table S2.**
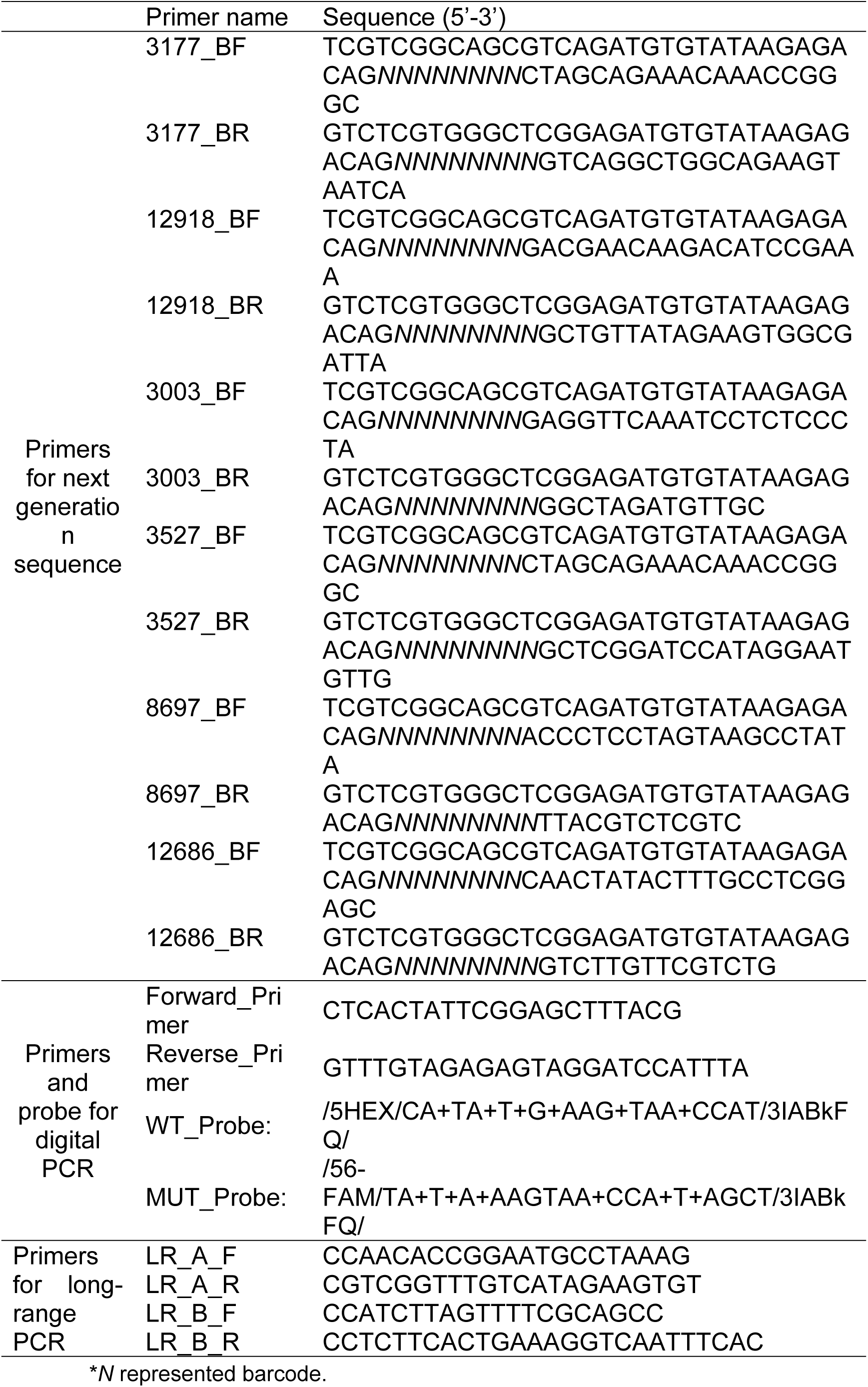
Primers used for sanger sequence, next generation sequence, digital PCR and long-range PCR.

## References

1. Anderson, S., Bankier, A.T., Barrell, B.G., de Bruijn, M.H.L., Coulson, A.R., Drouin, J., Eperon, I.C., Nierlich, D.P., Roe, B.A., Sanger, F., et al. (1981). Sequence and organization of the human mitochondrial genome. Nature 290, 457–465. 10.1038/290457a0.

2. Stewart, J.B., and Larsson, N.G. (2014). Keeping mtDNA in shape between generations. PLoS Genet 10, e1004670. 10.1371/journal.pgen.1004670.

3. Howe, D.K., and Denver, D.R. (2008). Muller’s Ratchet and compensatory mutation in Caenorhabditis briggsae mitochondrial genome evolution. BMC Evol Biol 8, 62. 10.1186/1471-2148-8-62.

4. Muller, H.J. (1964). The Relation of Recombination to Mutational Advance. Mutat Res 106, 2–9. 10.1016/0027-5107(64)90047-8.

5. Stewart, J.B., and Chinnery, P.F. (2015). The dynamics of mitochondrial DNA heteroplasmy: implications for human health and disease. Nature Reviews Genetics 16, 530–542. 10.1038/nrg3966.

6. Ng, Y.S., Bindoff, L.A., Gorman, G.S., Klopstock, T., Kornblum, C., Mancuso, M., McFarland, R., Sue, C.M., Suomalainen, A., Taylor, R.W., et al. (2021). Mitochondrial disease in adults: recent advances and future promise. The Lancet Neurology 20, 573–584. 10.1016/S1474-4422(21)00098-3.

7. Stewart, J.B., Freyer, C., Elson, J.L., Wredenberg, A., Cansu, Z., Trifunovic, A., and Larsson, N.G. (2008). Strong purifying selection in transmission of mammalian mitochondrial DNA. PLoS Biol 6, e10. 10.1371/journal.pbio.0060010.

8. Fan, W., Waymire, K.G., Narula, N., Li, P., Rocher, C., Coskun, P.E., Vannan, M.A., Narula, J., Macgregor, G.R., and Wallace, D.C. (2008). A mouse model of mitochondrial disease reveals germline selection against severe mtDNA mutations. Science 319, 958–962. 10.1126/science.1147786.

9. Lieber, T., Jeedigunta, S.P., Palozzi, J.M., Lehmann, R., and Hurd, T.R. (2019). Mitochondrial fragmentation drives selective removal of deleterious mtDNA in the germline. Nature 570, 380–384. 10.1038/s41586-019-1213-4.

10. Chen, Z., Wang, Z.H., Zhang, G., Bleck, C.K.E., Chung, D.J., Madison, G.P., Lindberg, E., Combs, C., Balaban, R.S., and Xu, H. (2020). Mitochondrial DNA segregation and replication restrict the transmission of detrimental mutation. J Cell Biol 219. 10.1083/jcb.201905160.

11. Chiang, A.C., McCartney, E., O’Farrell, P.H., and Ma, H. (2019). A Genome-wide Screen Reveals that Reducing Mitochondrial DNA Polymerase Can Promote Elimination of Deleterious Mitochondrial Mutations. Curr Biol 29, 4330–4336.e4333. 10.1016/j.cub.2019.10.060.

12. Ma, H., Xu, H., and O’Farrell, P.H. (2014). Transmission of mitochondrial mutations and action of purifying selection in Drosophila melanogaster. Nat Genet 46, 393–397. 10.1038/ng.2919.

13. Meshnik, L., Bar-Yaacov, D., Kasztan, D., Neiger, T., Cohen, T., Kishner, M., Valenci, I., Dadon, S., Klein, C.J., Vance, J.M., et al. (2022). Mutant C. elegans mitofusin leads to selective removal of mtDNA heteroplasmic deletions across generations to maintain fitness. BMC Biol 20, 40. 10.1186/s12915-022-01241-2.

14. 14. Li, M., Rothwell, R., Vermaat, M., Wachsmuth, M., Schröder, R., Laros, J.F., van Oven, M., de Bakker, P.I., Bovenberg, J.A., van Duijn, C.M., et al. (2016). Transmission of human mtDNA heteroplasmy in the Genome of the Netherlands families: support for a variable-size bottleneck. Genome Res 26, 417–426. 10.1101/gr.203216.115.

15. Wei, W., Tuna, S., Keogh, M.J., Smith, K.R., Aitman, T.J., Beales, P.L., Bennett, D.L., Gale, D.P., Bitner-Glindzicz, M.A.K., Black, G.C., et al. (2019). Germline selection shapes human mitochondrial DNA diversity. Science 364. 10.1126/science.aau6520.

16. Zaidi, A.A., Wilton, P.R., Su, M.S., Paul, I.M., Arbeithuber, B., Anthony, K., Nekrutenko, A., Nielsen, R., and Makova, K.D. (2019). Bottleneck and selection in the germline and maternal age influence transmission of mitochondrial DNA in human pedigrees. Proc Natl Acad Sci U S A 116, 25172–25178. 10.1073/pnas.1906331116.

17. Gupta, R., Kanai, M., Durham, T.J., Tsuo, K., McCoy, J.G., Kotrys, A.V., Zhou, W., Chinnery, P.F., Karczewski, K.J., Calvo, S.E., et al. (2023). Nuclear genetic control of mtDNA copy number and heteroplasmy in humans. Nature 620, 839–848. 10.1038/s41586-023-06426-5.

18. Cao, L., Shitara, H., Horii, T., Nagao, Y., Imai, H., Abe, K., Hara, T., Hayashi, J., and Yonekawa, H. (2007). The mitochondrial bottleneck occurs without reduction of mtDNA content in female mouse germ cells. Nat Genet 39, 386–390. 10.1038/ng1970.

19. Shoubridge, E.A., and Wai, T. (2007). Mitochondrial DNA and the mammalian oocyte. Curr Top Dev Biol 77, 87–111. 10.1016/s0070-2153(06)77004-1.

20. 20. Mok, B.Y., de Moraes, M.H., Zeng, J., Bosch, D.E., Kotrys, A.V., Raguram, A., Hsu, F., Radey, M.C., Peterson, S.B., Mootha, V.K., et al. (2020). A bacterial cytidine deaminase toxin enables CRISPR-free mitochondrial base editing. Nature 583, 631–637. 10.1038/s41586-020-2477-4.

21. 21. Yu-Wai-Man, P., and Chinnery, P.F. (1993). Leber Hereditary Optic Neuropathy. In GeneReviews(®), M.P. Adam, J. Feldman, G.M. Mirzaa, R.A. Pagon, S.E. Wallace, L.J.H. Bean, K.W. Gripp, and A. Amemiya, eds. (University of Washington, Seattle Copyright © 1993-2023, University of Washington, Seattle. GeneReviews is a registered trademark of the University of Washington, Seattle. All rights reserved.).

22. Morohaku, K., Tanimoto, R., Sasaki, K., Kawahara-Miki, R., Kono, T., Hayashi, K., Hirao, Y., and Obata, Y. (2016). Complete in vitro generation of fertile oocytes from mouse primordial germ cells. Proc Natl Acad Sci U S A 113, 9021–9026. 10.1073/pnas.1603817113.

23. Ota, S., Ikeda, S., Takashima, T., and Obata, Y. (2021). Optimal conditions for mouse follicle culture. J Reprod Dev 67, 327–331. 10.1262/jrd.2021-091.

24. Cheng, Y., Kawamura, K., Takae, S., Deguchi, M., Yang, Q., Kuo, C., and Hsueh, A.J. (2013). Oocyte-derived R-spondin2 promotes ovarian follicle development. Faseb j 27, 2175–2184. 10.1096/fj.12-223412.

25. Sato, Y., Cheng, Y., Kawamura, K., Takae, S., and Hsueh, A.J. (2012). C-type natriuretic peptide stimulates ovarian follicle development. Mol Endocrinol 26, 1158–1166. 10.1210/me.2012-1027.

26. Kawamura, K., Kawamura, N., Mulders, S.M., Sollewijn Gelpke, M.D., and Hsueh, A.J. (2005). Ovarian brain-derived neurotrophic factor (BDNF) promotes the development of oocytes into preimplantation embryos. Proc Natl Acad Sci U S A 102, 9206–9211. 10.1073/pnas.0502442102.

27. O’Brien, M.J., Pendola, J.K., and Eppig, J.J. (2003). A revised protocol for in vitro development of mouse oocytes from primordial follicles dramatically improves their developmental competence. Biol Reprod 68, 1682–1686. 10.1095/biolreprod.102.013029.

28. Floros, V.I., Pyle, A., Dietmann, S., Wei, W., Tang, W.C.W., Irie, N., Payne, B., Capalbo, A., Noli, L., Coxhead, J., et al. (2018). Segregation of mitochondrial DNA heteroplasmy through a developmental genetic bottleneck in human embryos. Nature Cell Biology 20, 144–151. 10.1038/s41556-017-0017-8.

29. Freyer, C., Cree, L.M., Mourier, A., Stewart, J.B., Koolmeister, C., Milenkovic, D., Wai, T., Floros, V.I., Hagström, E., Chatzidaki, E.E., et al. (2012). Variation in germline mtDNA heteroplasmy is determined prenatally but modified during subsequent transmission. Nature Genetics 44, 1282–1285. 10.1038/ng.2427.

30. Kauppila, J.H.K., Baines, H.L., Bratic, A., Simard, M.L., Freyer, C., Mourier, A., Stamp, C., Filograna, R., Larsson, N.G., Greaves, L.C., and Stewart, J.B. (2016). A Phenotype-Driven Approach to Generate Mouse Models with Pathogenic mtDNA Mutations Causing Mitochondrial Disease. Cell Rep 16, 2980–2990. 10.1016/j.celrep.2016.08.037.

31. Zhang, H., Esposito, M., Pezet, M.G., Aryaman, J., Wei, W., Klimm, F., Calabrese, C., Burr, S.P., Macabelli, C.H., Viscomi, C., et al. (2021). Mitochondrial DNA heteroplasmy is modulated during oocyte development propagating mutation transmission. Sci Adv 7, eabi5657. 10.1126/sciadv.abi5657.

32. Kidere, D., Zayakin, P., Livcane, D., Makrecka-Kuka, M., Stavusis, J., Lace, B., Lin, T.K., Liou, C.W., and Inashkina, I. (2023). Impact of the m.13513G>A Variant on the Functions of the OXPHOS System and Cell Retrograde Signaling. Curr Issues Mol Biol 45, 1794–1809. 10.3390/cimb45030115.

33. Wang, Z., Qi, X.K., Yao, S., Chen, B., Luan, X., Zhang, W., Han, M., and Yuan, Y. (2010). Phenotypic patterns of MELAS/LS overlap syndrome associated with m.13513G>A mutation, and neuropathological findings in one autopsy case. Neuropathology 30, 606–614. 10.1111/j.1440-1789.2010.01115.x.

34. Hill, J.H., Chen, Z., and Xu, H. (2014). Selective propagation of functional mitochondrial DNA during oogenesis restricts the transmission of a deleterious mitochondrial variant. Nat Genet 46, 389–392. 10.1038/ng.2920.

35. Jeedigunta, S.P., Minenkova, A.V., Palozzi, J.M., and Hurd, T.R. (2021). Avoiding Extinction: Recent Advances in Understanding Mechanisms of Mitochondrial DNA Purifying Selection in the Germline. Annu Rev Genomics Hum Genet 22, 55–80. 10.1146/annurev-genom-121420-081805.

36. 36. Ma, H., Hayama, T., Van Dyken, C., Darby, H., Koski, A., Lee, Y., Gutierrez, N.M., Yamada, S., Li, Y., Andrews, M., et al. (2020). Deleterious mtDNA mutations are common in mature oocytes. Biol Reprod 102, 607–619. 10.1093/biolre/ioz202.

37. Franco, M., Pickett, S.J., Fleischmann, Z., Khrapko, M., Cote-L’Heureux, A., Aidlen, D., Stein, D., Markuzon, N., Popadin, K., Braverman, M., et al. (2022). Dynamics of the most common pathogenic mtDNA variant m.3243A > G demonstrate frequency-dependency in blood and positive selection in the germline. Hum Mol Genet 31, 4075–4086. 10.1093/hmg/ddac149.

38. Cote-LHeureux, A., Fleischmann, Z., Franco, M., Chen, Z., Khrapko, M., Vyshedskiy, B., Braverman, M., Popadin, K., Pickett, S., Woods, D.C., et al. (2023). A novel, wave-shaped profile of germline selection of pathogenic mtDNA mutations is discovered by bypassing a classical statistical bias. Genomics preprint. 10.1101/2023.11.21.568140.

39. Stewart, J.B., Freyer, C., Elson, J.L., and Larsson, N.G. (2008). Purifying selection of mtDNA and its implications for understanding evolution and mitochondrial disease. Nat Rev Genet 9, 657–662. 10.1038/nrg2396.

40. Hsueh, A.J., Billig, H., and Tsafriri, A. (1994). Ovarian follicle atresia: a hormonally controlled apoptotic process. Endocr Rev 15, 707–724. 10.1210/edrv-15-6-707.

41. Adashi, E.Y., and Cohen, I.G. (2018). Preventing Mitochondrial Diseases: Embryo-Sparing Donor-Independent Options. Trends Mol Med 24, 449–457. 10.1016/j.molmed.2018.03.002.

42. Zhang, Y., Wang, Z.H., Liu, Y., Chen, Y., Sun, N., Gucek, M., Zhang, F., and Xu, H. (2019). PINK1 Inhibits Local Protein Synthesis to Limit Transmission of Deleterious Mitochondrial DNA Mutations. Mol Cell 73, 1127–1137.e1125. 10.1016/j.molcel.2019.01.013.

43. Kremer, L.S., Bozhilova, L.V., Rubalcava-Gracia, D., Filograna, R., Upadhyay, M., Koolmeister, C., Chinnery, P.F., and Larsson, N.G. (2023). A role for BCL2L13 and autophagy in germline purifying selection of mtDNA. PLoS Genet 19, e1010573. 10.1371/journal.pgen.1010573.

44. Ru, Y., Deng, X., Chen, J., Zhang, L., Xu, Z., Lv, Q., Long, S., Huang, Z., Kong, M., Guo, J., and Jiang, M. (2024). Maternal age enhances purifying selection on pathogenic mutations in complex I genes of mammalian mtDNA. Nat Aging 4, 1211–1230. 10.1038/s43587-024-00672-6.

